# The properties of human disease mutations at protein interfaces

**DOI:** 10.1101/2021.08.20.457107

**Authors:** Benjamin J. Livesey, Joseph A. Marsh

## Abstract

The assembly of proteins into complexes and interactions with other biomolecules are often vital for their biological function. While it is known that mutations at protein interfaces have a high potential to be damaging and cause human genetic disease, there has been relatively little consideration for how this varies between different types of interfaces. Here we investigate the properties of human pathogenic and putatively benign missense variants at homomeric (isologous and heterologous), heteromeric, DNA, RNA and other ligand interfaces, and at different regions with respect to those interfaces. We find that different types of interfaces vary greatly in their propensity to be associated with pathogenic mutations, with homomeric heterologous and DNA interfaces being particularly enriched in disease. We also find that residues that do not directly participate in an interface, but are close in 3D space, also show a significant disease enrichment. Finally, we show that mutations at different types of interfaces tend to have distinct property changes when undergoing amino acid substitutions associated with disease, and that this is linked to substantial variability in their identification by computational variant effect predictors.

## Introduction

Single nucleotide variants (SNVs) are the most common type of human genetic variation (1). While many SNVs in protein-coding regions of the genome are associated with human disease, the vast majority have no noticeable clinical impact (2). We have still seen only a fraction of the possible coding SNVs in genetic sequencing studies and most possible variants remain completely unannotated. Distinguishing novel disease-causing SNVs from those that are benign is therefore a major ongoing challenge for the field of bioinformatics.

Protein misfolding and destabilization have long been held as primary mechanisms by which mutations cause disease (3), and it is well established that pathogenic missense mutations are enriched within interior residues of proteins relative to the surface (4–6). In the last decade, there has also been increasing recognition of the importance of protein interfaces as hubs for disease-associated mutants (7–9). A mutation at an interface can act to destabilize the interface, disrupt the binding of the partner, increase, reduce or alter the affinity of the partner or stabilize the interface (10), an association which has been shown to be highly associated with disease states (11). Experimental work has supported the idea that a large fraction of pathogenic alleles do not destabilise protein folding but instead perturb protein interactions (12).

The separation of interface residues into core and rim regions can be useful for understanding the mechanisms of pathogenesis at protein interfaces. Core regions are those buried within the interface and contain the highest enrichment of pathogenic variants (11,13,14). Rim regions are located around the edge of the interface and are more exposed to solvent. Core and rim residues show differences in both physiochemical properties and conservation (11,15). There is also evidence that rim residue mutations may be more impactful in smaller complexes (16).

It was noted some time ago, that mutations in specific “hot spot” residues within interfaces can greatly alter the free energy of binding (17) through disruption of non-covalent interactions (18). It has since been shown that mutations of these hot spot residues are enriched for disease mutations relative to the rest of the interface (11,19,20). Hot spots are preferentially found within the interface core, although those located in the rim remain enriched for pathogenic variants (11).

Interface mutations also appear to be particularly important in the context of cancer (21–23). Those proteins impacted also tended to be hub proteins within the human interactome. In p53 and other proteins, it was shown that patient survival was correlated to the specific interactions that were perturbed (21,24). Mutations in interfaces allow for specific interactions to be abolished and altered, while the protein retains some activity (25). This ‘edgetic’ perturbation allows cancer to re-wire the normal interactome with less risk of full collapse that may result from full protein destabilisation. A similar pattern of interaction destabilizing mutants has also been observed for developmental disorders such as autism (26).

To improve the prediction of protein variant effects, it is important to first better understand the molecular mechanisms underlying damaging mutations and how these are related to amino acid properties. Several studies have shown that tryptophan, tyrosine and arginine mutations in interfaces are overrepresented among disease-causing mutations (13,17). Different amino acid substitutions also cause disease at different rates depending on where they occur in the protein or interface (11). Drastic changes in physio-chemical properties at interfaces were observed in cancer-associated mutations (22). However, other studies have demonstrated that amino acid conservation is often a larger factor in predicting pathogenic mutations than the change in properties (27). As with the protein interior residues, conservation remains an important factor for understanding the pathogenicity of mutations at protein interfaces (11,15).

One aspect that has not received much consideration in previous studies of protein interfaces, is the nature and orientation of the interactors. Protein-protein interfaces are most easily divided by considering homomeric and heteromeric interfaces separately. Homomers can be further divided into isologous (head-to-head) and heterologous (head-to-tail) interactors. Proteins also interface with numerous non-protein biomolecules such as DNA and RNA. In this study, we have taken interface type into consideration, in an attempt to better understand how the structural and sequence properties of different types of human protein interfaces are related to the effects of mutations. We investigate the enrichment of pathogenic mutations within different regions of the proteins and types of interface. We also investigate the property changes caused by mutations, and how these differ between protein regions. Finally, we relate how the findings of this study have implications for the future of protein variant effect prediction in protein-protein and protein-biomolecule interfaces.

## Results and discussion

### Enrichment of disease mutations at different types of protein interfaces

To represent pathogenic variants in our study, we used missense variants from the ClinVar (28) database, including only those annotated as ‘pathogenic’ or ‘likely pathogenic’. We also used the gnomAD v2.1 (29) database of variants observed in the human population, excluding those also present in the ClinVar dataset. We refer to these variants as ‘putatively benign’, since while gnomAD excludes individuals with severe paediatric disease, recessive and low-penetrance variants are certainly present in the dataset to some extent. We considered only variants that could be mapped to residues in published PDB structures; this resulted in a total of 495,076 putatively benign missense variants in 4033 genes, and 19,175 (likely) pathogenic missense variants in 1754 genes.

First, we looked at how the locations of missense variants within protein and protein complex structures are related to enrichment in disease. We used solvent accessible surface area (SASA) to classify residues in PDB structures of human proteins as surface, interior or interface (Fig 1A). Surface residues have 25% or more of their surface area (30) exposed to solvent, while interior residues have less than this. Interface residues undergo a change in SASA between the monomeric and complexed forms of the structure, meaning they participate directly in the interface.

**Figure 1:**
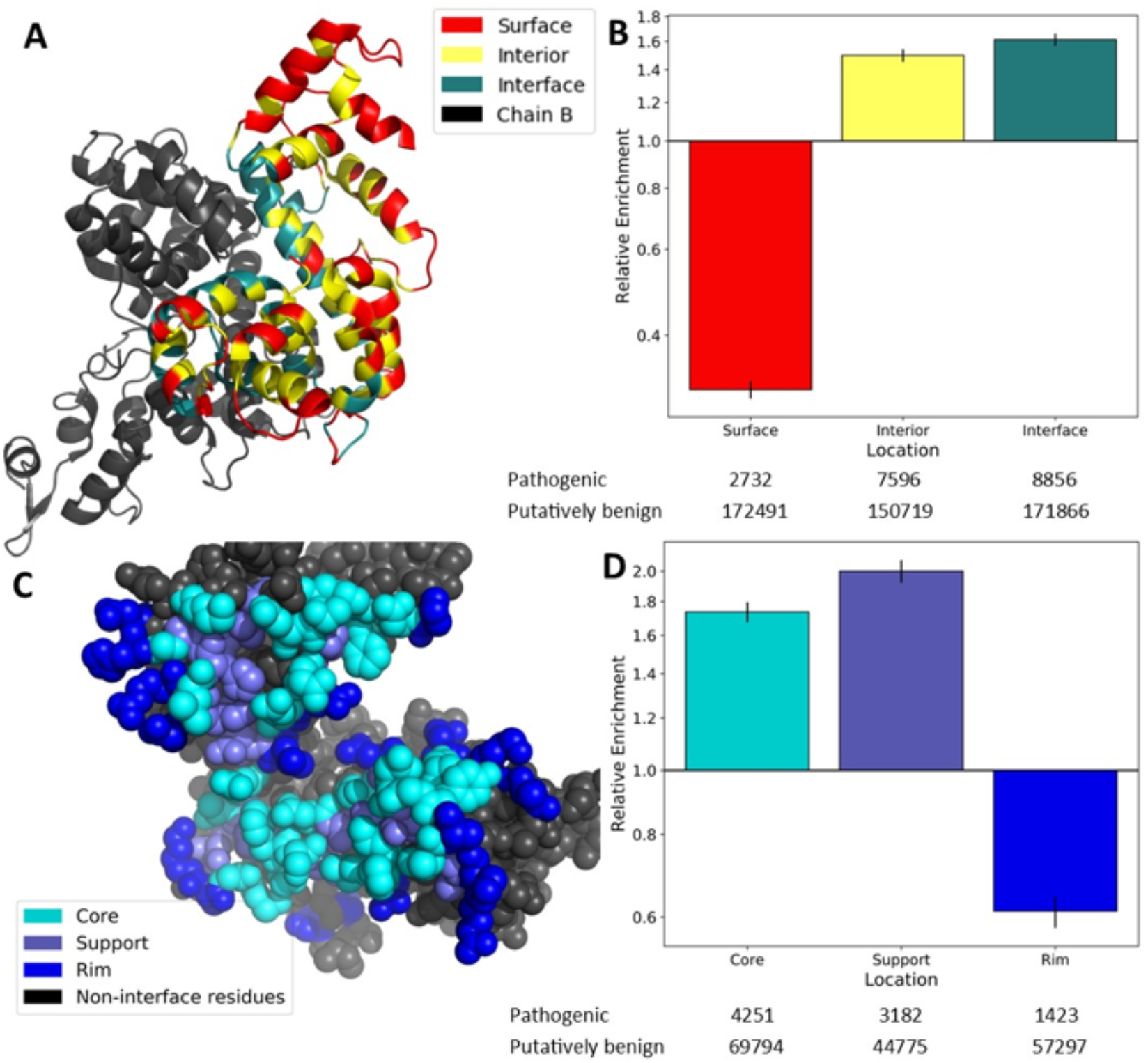
The anatomy of protein interfaces and the enrichment of pathogenic mutations. A) Ribbon representation of the structure of a homodimeric mitochondrial aspartate/glutamate carrier protein (69) (PDB ID: 4P60), highlighting surface, interior and interface residues. B) Relative enrichment (odds ratio) of pathogenic mutations within protein surface, interior and interface locations. C) Sphere representation of the 4P60 dimer homomeric interface on one chain, highlighting core, support and rim residues. D) Relative enrichment of pathogenic mutations within interface core, support and rim regions. All error bars represent 95% confidence intervals. Numbers of pathogenic and putatively benign mutations within each location dataset are shown below the plots.

By calculating the odds ratio of pathogenic to putatively benign variants at each location, we find that mutations on the protein surface are highly depleted of pathogenic variants (Fig 1B) (0.31 times, p<2.225×10^−308^), while interior positions are enriched (1.50 times, p=1.06×10^−153^). This interior enrichment differs somewhat from a previous estimate (2.66 times), but that study used different mutation datasets and a stricter definition of buried residues (<1% exposed surface area) (31). Other studies which used a similar definition of interior residues to us obtained similar enrichment values (32). Interestingly, we find a greater enrichment of disease mutations than even interior positions within protein interfaces of 1.61 times (p=1.83×10^−225^). Another study (11), found that disease enrichment in interface residues was intermediate to interior and surface residues. This study also used a stricter definition for interior residues (7% SASA cutoff) as well as a definition for interface residues based on residue proximity rather than SASA changes, potentially accounting for the difference.

Next, we split the interface residues into three regions – core, rim and support – as previously defined (33). Core interface residues are on the surface of the unbound subunit (≥ 25% of their surface area exposed in the monomer), but are buried in the bound complex, suggesting that they are likely to be crucial for the interaction. Support residues are buried in the unbound subunit (<25% of surface area exposed), suggesting that their main role may be in stabilising intramolecular structure, but they may also participate in the interaction. Rim residues are on the surface of both the monomer and full complex (≥25% surface area exposed in both), suggesting that they may be less important to both protein stability and the interaction. An example of these classifications is shown in Fig 1C.

The distribution of pathogenic variants within the different interface regions is not uniform. Both support and core residues show a strong enrichment in pathogenic variants (2.00 times and 1.74 times respectively). In contrast, rim residues resemble surface residues in that they show a depletion of pathogenic mutants (0.61 times, p=1.68×10^−78^). These results largely agree with previous studies of interface mutations where variants buried within protein interfaces demonstrate a greater tendency to cause disease than those on the interface rim (8,11). Due to the fact that rim residues are most similar to surface residues in terms of their lack of disease enrichment, for subsequent analyses, we have grouped rim residues with other surface residues rather than interfaces.

While previous work has clearly demonstrated the importance of protein interfaces for understanding disease mutations, there has been relatively little consideration of the different types of protein interfaces. Therefore, we further classified protein interfaces based upon the types of interactions they make (Fig 2A). Homomeric interfaces, formed on interaction between two copies of the same protein subunit, can be split up into those that are isologous (i.e. head-to-head or symmetric, involving the same surface patch on each subunit), and those that are heterologous (i.e. head-to-tail or asymmetric, involving different surface patches on each subunit) (34). We also separately considered heteromeric interfaces (formed between two distinct protein subunits), as well as DNA, RNA and other ligand-binding interfaces.

**Figure 2:**
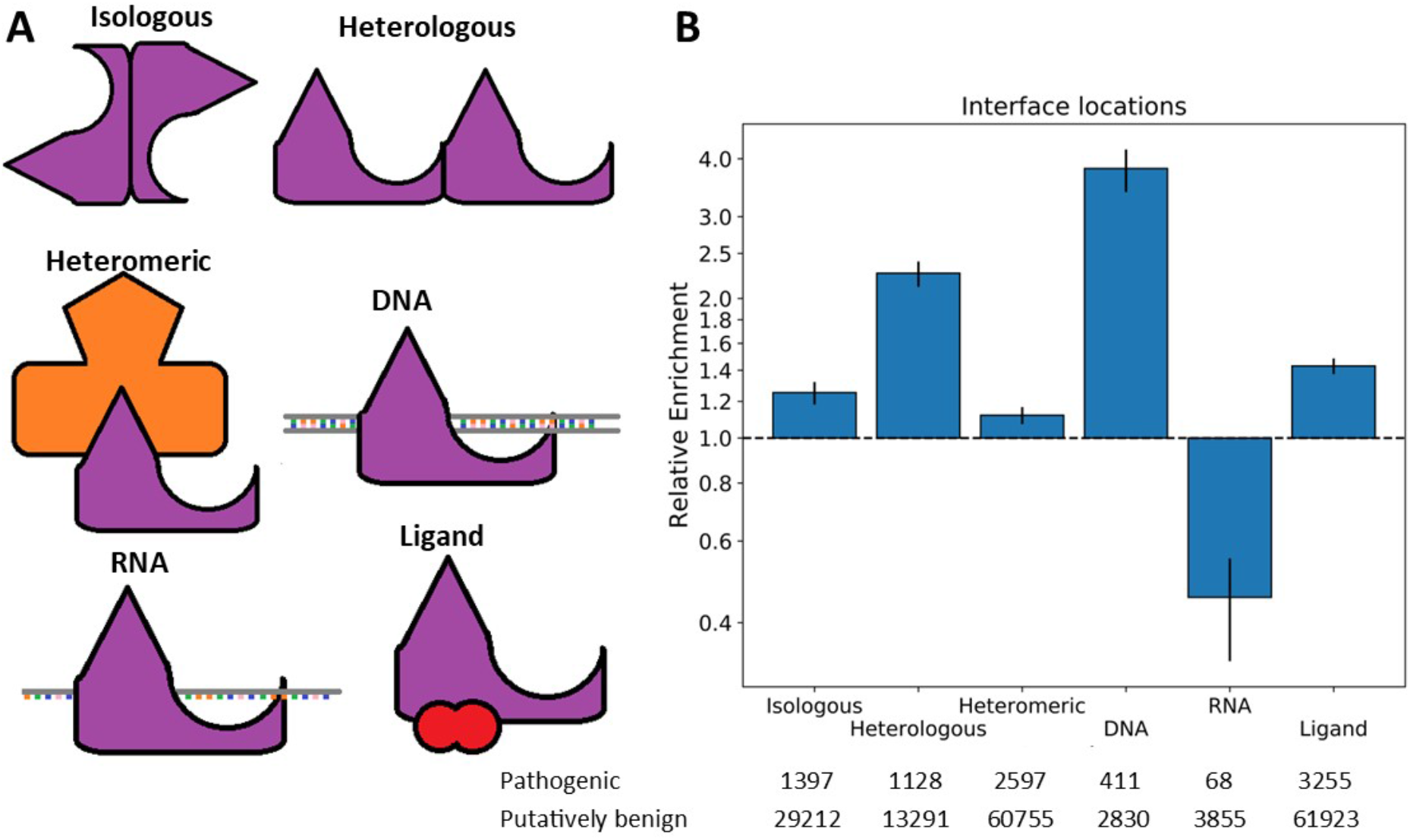
Comparison of the enrichment of pathogenic mutations at different protein interfaces. A) Demonstration of each protein interface types. B) Relative enrichment of pathogenic mutations within each interface type. Error bars represent 95% confidence intervals. Numbers of pathogenic and putatively benign mutations within each location dataset are shown.

Again, we calculated the odds ratios of pathogenic to benign variants for each interface type (Fig 2B). Strikingly, DNA interfaces showed the greatest enrichment of pathogenic variants (3.81 times, p=8.54×10^−102^), which we suspect may be at least partially due to the highly damaging effects of transcription factors with off-target effects (Williamson *et al*, 2019; Latchman, 1996). We also observed that heterologous interfaces were more disease-enriched (2.26 times, p=1.26×10^−118^) than either isologous interfaces (1.25 times, p=1.29×10^−14^) or heteromeric interfaces (1.12 times, p=2.15×10^−7^). Surprisingly, RNA interface residues were in fact depleted in disease mutations (0.45 times, p=2.37×10^−13^). However, after excluding ribosomal proteins from the analysis, the enrichment becomes 1.04 (p=0.71). One possible explanation for this is that ribosomal proteins are so highly conserved that diverse impactful variants are simply not observed in the population due to embryonic lethality.

The fact that pathogenic mutations are enriched in homomeric heterologous compared to homomeric isologous interfaces is very interesting. Homomeric interface type is closely related to complex symmetry, with symmetric dimers (*C*_2_) having isologous interfaces, complexes with higher order cyclic symmetry (*C*_n; n>2_) having heterologous interfaces, dihedral complexes (*D*_n_) having isologous and sometimes heterologous interfaces, and complexes with helical (*H*) symmetry having exclusively heterologous interfaces (37). Therefore, it is possible that this trend is driven by a disease association with complex symmetry, as has previously been suggested (38). We therefore investigated the enrichment of disease mutations within complexes of different symmetry (Fig 3A). Our results indicate that monomers and symmetric dimers show a slight depletion in pathogenic mutants. Larger cyclic, dihedral and helical complexes on the other hand showed a slight enrichment. One possible explanation for this could be that that they simply have relatively fewer surface residues and more interface residues. However, we observe that interior and interface residues in cyclic and dihedral complexes show a greater disease enrichment than in monomers and symmetric dimers (Fig 3B). While interior residues in helical complexes are actually depleted in disease mutations, it is important to note the small number of helical complexes in our dataset and the corresponding large range covered by the confidence interval.

**Figure 3:**
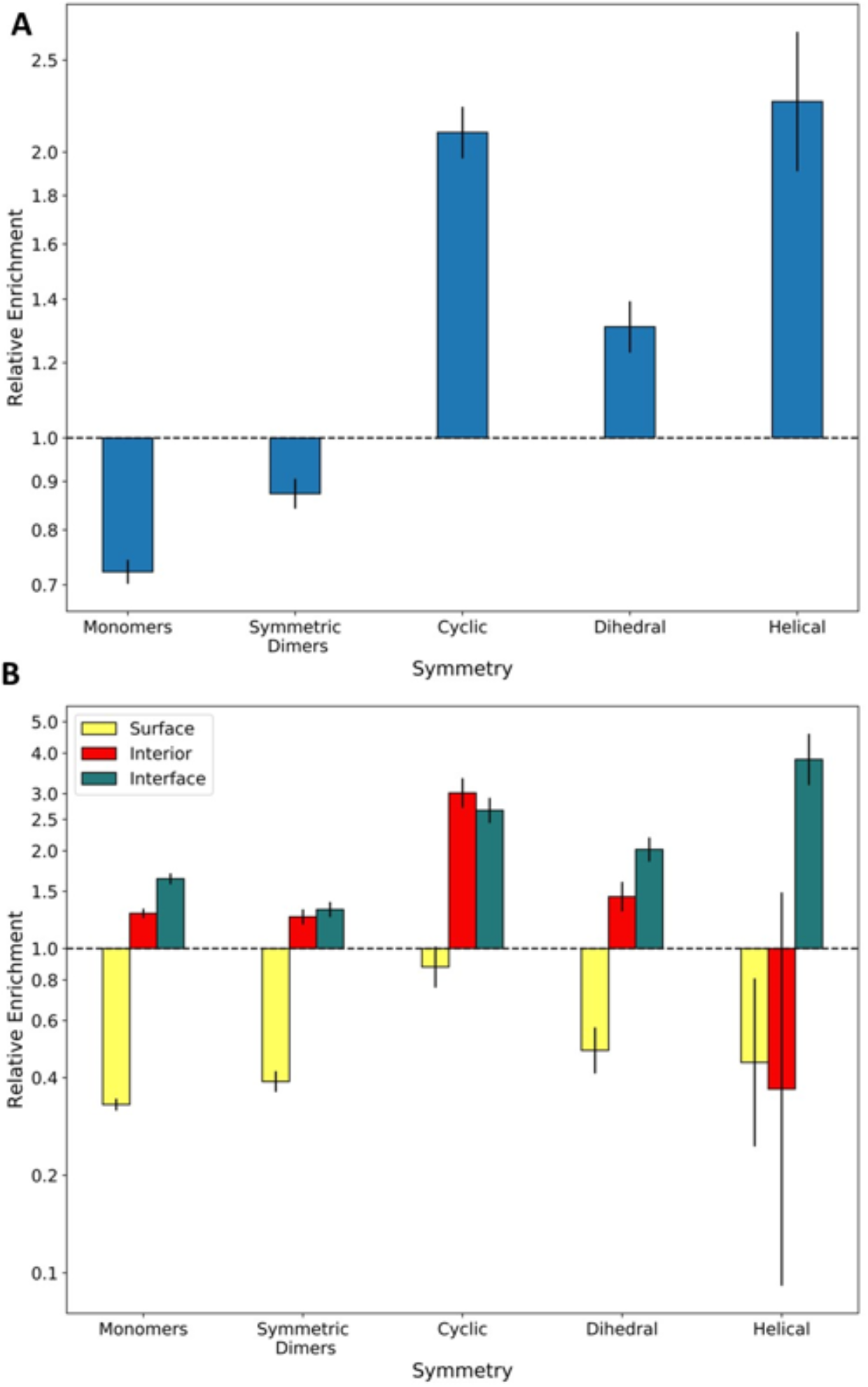
The enrichment of pathogenic mutations within protein complexes of different symmetries. A) Relative enrichment of pathogenic mutations within proteins of each symmetry type. Symmetric dimers (C_2_) are considered separately from higher-order cyclic (C_n; n>2_) complexes. Complexes with cubic symmetries are not included due to lack of representation in the data. B) Enrichment of pathogenic mutations in the surface, interior and interface of each symmetry type relative to the full dataset. All error bars represent 95% confidence intervals.

It is notable that all of the enriched complex types can harbour heterologous homomeric interfaces. The observed enrichment may also be related to functional associations of the different symmetry types, *e*.*g*. dihedral complexes are often enzymes while cyclic complexes include many transmembrane channels (39). To investigate if a high proportion of membrane proteins with heterologous interfaces were biasing our analysis, we repeated the calculations while excluding all proteins tagged with the keyword ‘membrane’ (KW-0472) in UniProt (Fig S1). Heterologous interfaces remained more enriched for pathogenic variants (1.87 times, p=6.93×10^−33^) than either isologous or heteromeric interfaces.

### Enrichment of disease mutations in proximity to interfaces

Our observations show that the support residues of protein interfaces, which are most buried intramolecularly within the protein structure, clearly have a greater enrichment of disease mutations than the interface core, or interior residues. This is most likely due to the combined effect of support mutations destabilising both folding and intermolecular interactions. We therefore hypothesised that other interior residues in proximity to the interface would show similar levels of disease enrichment, despite not participating in the interface directly.

To address this, we calculated the proximity of residues to the nearest interface and compared this to their disease enrichment. We utilized the interface centroid, which is defined as the mean of the X, Y and Z coordinates of all residue alpha-carbons in the interface (Fig 4A). We generated 30 bins containing identical numbers of variants at increasing distance from the nearest interface centroid, and identified a strong correlation (−0.89 Spearman’s ρ) between this distance and the enrichment of disease mutations within the bins (Fig 4B). Notably, a greater correlation was found (−0.73) when we consider interior residues alone (Fig 4C), than when we consider surface residues alone (−0.63) (Fig 4D). Interior residues in close proximity to the nearest interface centroid can be considered as pseudo-support residues, directly influencing the buried residues in the interface. Our results suggest that this effect does not terminate directly adjacent to the interface and instead propagates further through the structure. The effect of interface proximity on disease-risk does not appear to influence the protein surface as much as the interior, possibly due to the lower density of intramolecular contacts available for propagating mutation effects.

**Figure 4:**
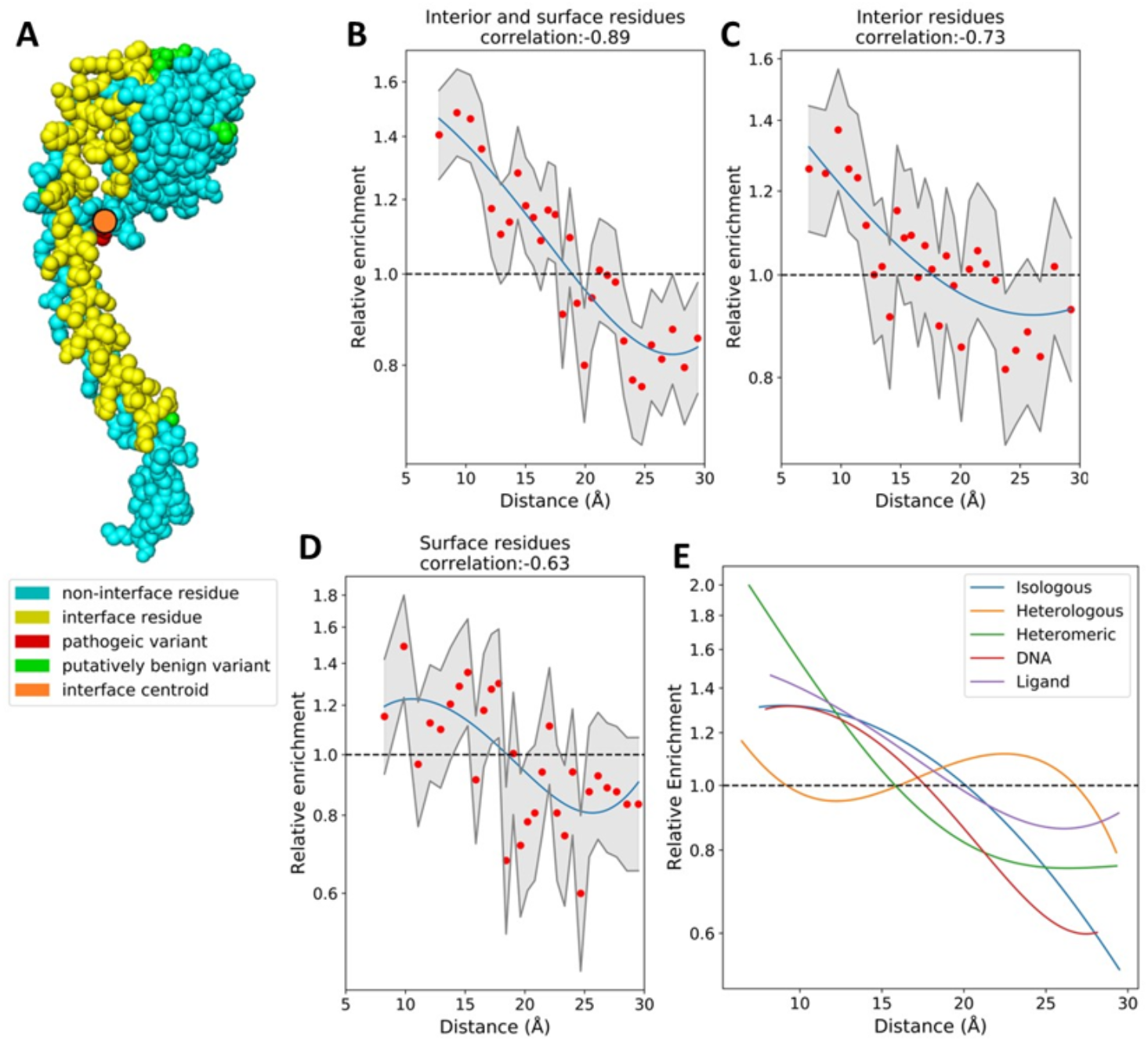
The relationship between interface proximity and disease enrichment. A) Sphere representation of a protein with interface residues highlighted in yellow. The interface centroid is an orange circle. Putatively benign mutations present in gnomAD are shown in green, while pathogenic ClinVar mutations are in red. B-D) Log odds ratio of disease mutations at increasing distance from the nearest interface centroid. 95% confidence intervals are shown in grey, a univariate spline (blue) has been fit to the data. Charts are shown for surface and interior residues together and individually. E) Overlaid splines fit to odds ratio data of the different protein interface types (taking interior and surface residues together).

Allosteric residues may contribute to this effect, by being mutated directly, or by assisting the propagation of the effect to the interface. Allosteric residues are known to be highly conserved and have been used to explain the effects of several uncharacterised pathogenic mutants (40). Recently, two allosteric F-actin mutants, which are distal to the myosin interface, were found to impact activity at that interface (41); this is similar to the overall pattern we observe.

In addition to the interface centroid, we also investigated the minimum distance between a mutation and any interface residue, and the mean distance to all residues in the interface as interface proximity metrics (Fig S2). A comparable pattern is found regardless of the metric used; however, the correlations are greatest when using centroid distance. Centroid distance is closely related to average distance, with a Spearman’s correlations of 0.93 between the two metrics. Minimum distance is less closely related with correlations of 0.65 to centroid distance and 0.59 to average distance.

We found a slight difference in the propagation of disease enrichment in different interface types (Fig 4E). Both isologous and heterologous homomeric interfaces display a greater enrichment of disease mutations between 15 and 25 Å from the interface centroid than heteromeric interfaces do. We also find that disease enrichment in DNA interfaces drops off more rapidly than in protein interfaces. We speculate that this could be due to specificity-altering mutations in DNA interfaces needing to be closer to the interface surface, while those further back either completely abolish the interaction or have no effect. Ligand interfaces show a shallower curve than any of the other interface types, although it crosses the neutral-enrichment point at roughly the same location, perhaps reflecting their smaller interfaces, or the fact that many ligands in protein structures are not biologically important (*e*.*g*. crystallisation artefacts).

### Properties of amino acid substitutions at interfaces

Given that protein interfaces tend to involve different types of amino acids than protein interiors or surfaces (33,42), we wondered whether this could be reflected in differential propensities for pathogenic amino acid substitutions. We therefore calculated the residue-level disease enrichment for both the wild-type and mutant amino acids for different locations (Fig 5 and Fig S3).

**Figure 5:**
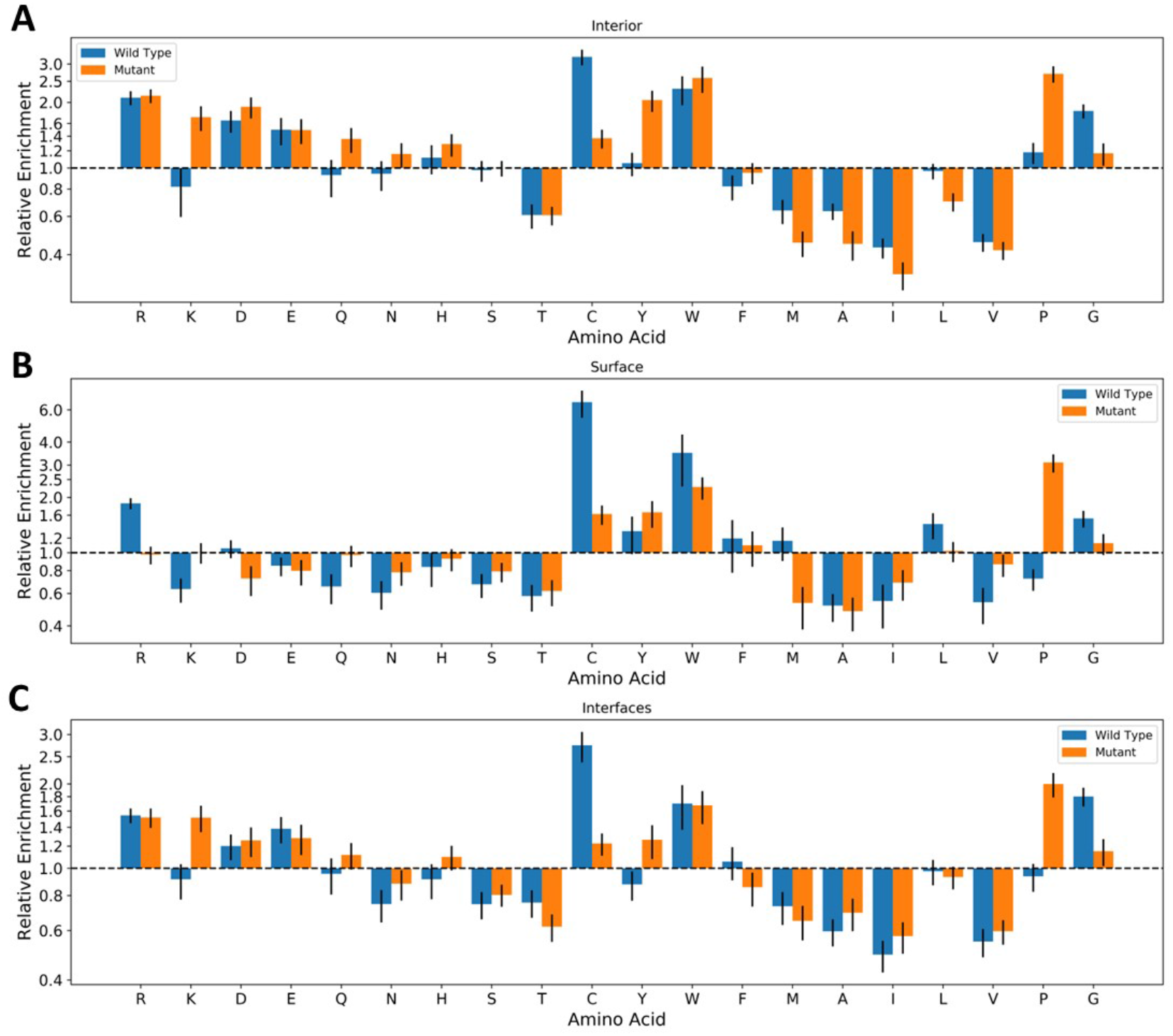
Enrichment of pathogenic variants at specific residues at different locations. The log odds ratio of pathogenic mutations associated with specific amino acid mutations at A) the protein interior, B) the protein surface and C) protein interfaces. Blue bars represent mutations *from* a specific amino acid, while orange bars represent mutations *to* a specific amino acid. All error bars represent 95% confidence intervals. Enrichments for different interface types are provided in Figure S3.

Across interior, surface and interface locations, the enrichments show broad similarities, with mutations from arginine, tryptophan, cysteine and glycine wild-type residues most likely to result in a pathogenic outcome, while mutations to tryptophan and proline were the most likely to be pathogenic. This largely agrees with previous findings (Vitkup et al, 2003). In surface residues, mutations from cysteine were more damaging than at interior locations. This is in-line with previous work, showing that thiol groups on protein surfaces play a role in regulating the cellular redox environment (43,44). Substitution of charged residues clearly results in more pathogenic mutants on the interior, while hydrophobic substitutions are more damaging on the surface with the exceptions of tryptophan and glycine. Aspartate mutations are differentially enriched (enriched in pathogenic mutants on the interior, but depleted on the surface), and clusters with other charged amino acids which are either less differentially enriched or neither enriched not depleted at surface locations.

Given that interface core and support residues are buried upon complex formation, we naturally can expect the mutational profile of interfaces to be similar to that of the protein interior. Comparing the relative enrichment of disease mutations in interior and interface mutations, we find that substitutions involving charged and polar amino acids as either the wild type or the mutant are more likely to be damaging for the interior. In contrast, small hydrophobic amino acid substitutions are more likely to be damaging within interfaces. These results imply that disruption of hydrophobic interactions is an important mechanism for pathogenesis in interfaces.

While investigating the mutational profiles of the different interface types we found that our analysis of the residue mutations lacked power due to sub-setting of the data across the large number of possible amino acid substitutions. To supplement this analysis, we downloaded 547 amino acid physical, biochemical and statistical properties from AAindex (45). By using amino acid properties, all mutations at a location can now contribute towards the results of our analysis. For each property, we calculated the difference between the wild type and mutant amino acids in our data sets. We then calculated the p-value and effect size of the mean difference between the datasets. Initially we found only small effect sizes (<0.3 Hedges’ g), however, taking the absolute difference between wild type and the mutant greatly increased the effect. This implies that the scale of deviation from the wild-type property is more important than the direction of the deviation for most pathogenic mutations. The top three properties by effect size per location are shown in Table 1. The full data is available in Table S1.

**Table 1:**
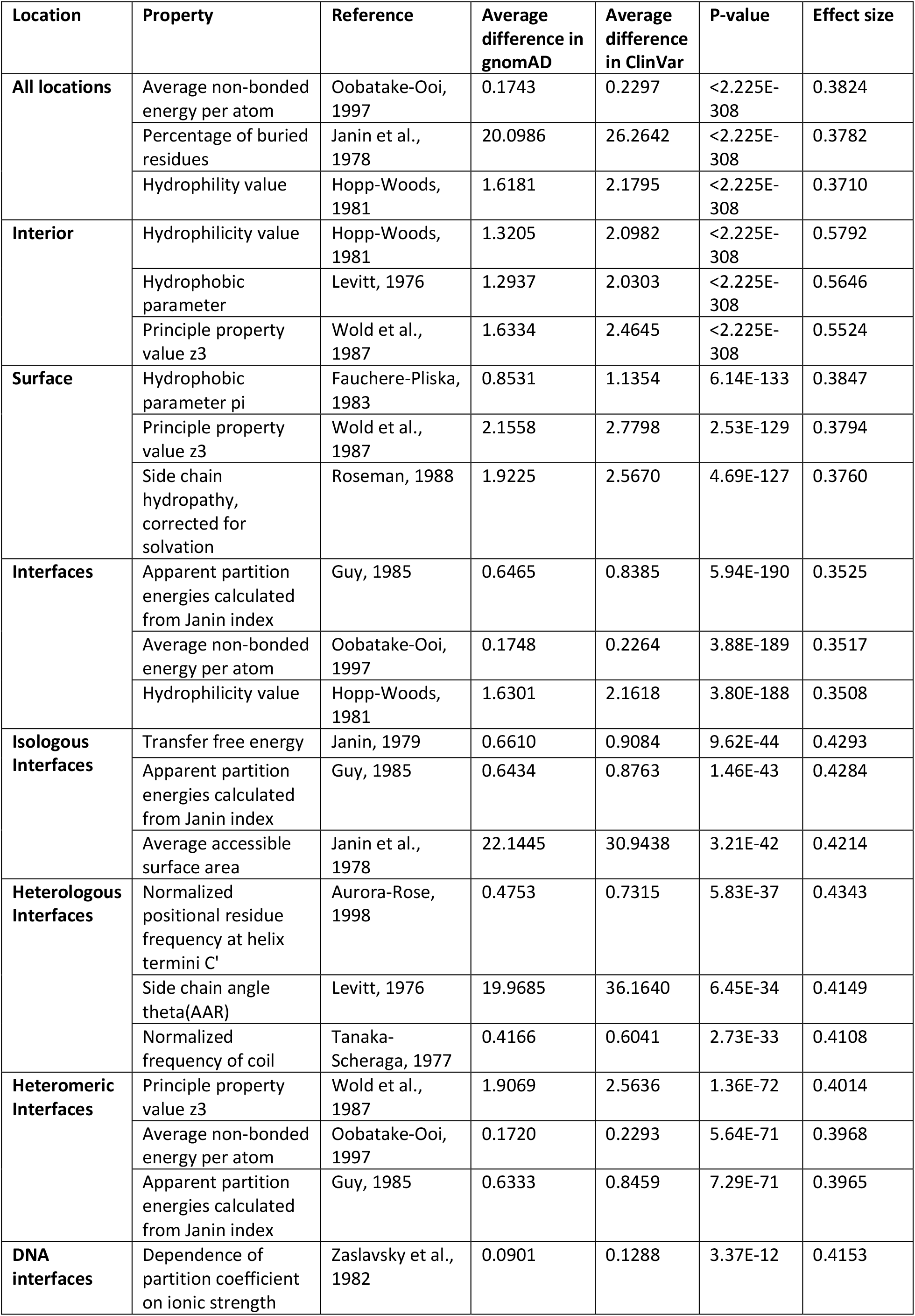
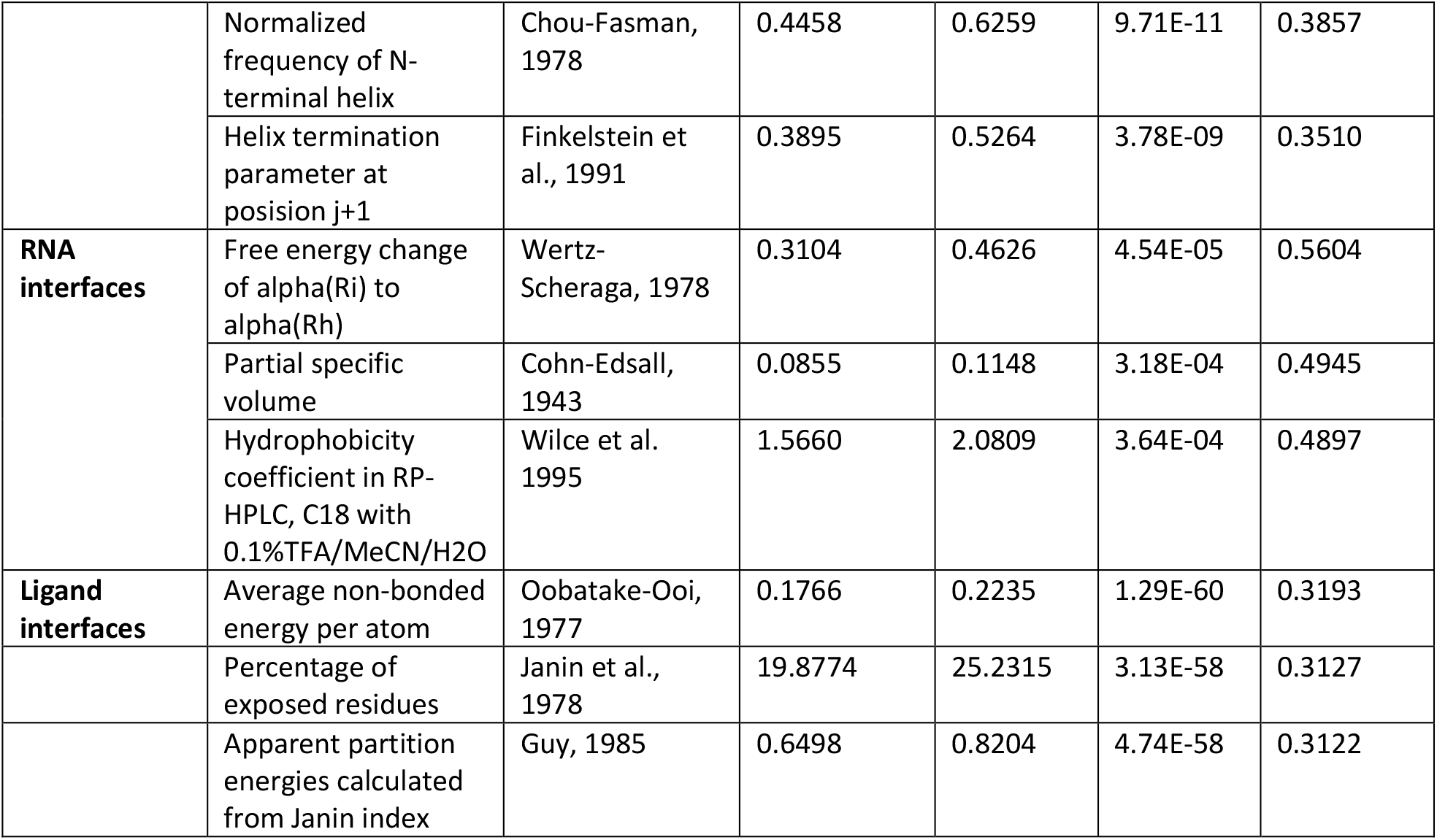
Top 3 property changes by effect size at different protein locations.

| Location | Property | Reference | Average difference in gnomAD | Average difference in ClinVar | P-value | Effect size |
| --- | --- | --- | --- | --- | --- | --- |
| **All locations** | Average non-bonded energy per atom | Oobatake-Ooi, 1997 | 0.1743 | 0.2297 | <2.225E-308 | 0.3824 |
|  | Percentage of buried residues | Janin et al., 1978 | 20.0986 | 26.2642 | <2.225E-308 | 0.3782 |
|  | Hydrophilicity value | Hopp-Woods, 1981 | 1.6181 | 2.1795 | <2.225E-308 | 0.3710 |
| **Interior** | Hydrophilicity value | Hopp-Woods, 1981 | 1.3205 | 2.0982 | <2.225E-308 | 0.5792 |
|  | Hydrophobic parameter | Levitt, 1976 | 1.2937 | 2.0303 | <2.225E-308 | 0.5646 |
|  | Principle property value z3 | Wold et al., 1987 | 1.6334 | 2.4645 | <2.225E-308 | 0.5524 |
| **Surface** | Hydrophobic parameter pi | Fauchere-Pliska, 1983 | 0.8531 | 1.1354 | 6.14E-133 | 0.3847 |
|  | Principle property value z3 | Wold et al., 1987 | 2.1558 | 2.7798 | 2.53E-129 | 0.3794 |
|  | Side chain hydrophathy, corrected for solvation | Roseman, 1988 | 1.9225 | 2.5670 | 4.69E-127 | 0.3760 |
| **Interfaces** | Apparent partition energies calculated from Janin index | Guy, 1985 | 0.6465 | 0.8385 | 5.94E-190 | 0.3525 |
|  | Average non-bonded energy per atom | Oobatake-Ooi, 1997 | 0.1748 | 0.2264 | 3.88E-189 | 0.3517 |
|  | Hydrophilicity value | Hopp-Woods, 1981 | 1.6301 | 2.1618 | 3.80E-188 | 0.3508 |
| **Isologous Interfaces** | Transfer free energy | Janin, 1979 | 0.6610 | 0.9084 | 9.62E-44 | 0.4293 |
|  | Apparent partition energies calculated from Janin index | Guy, 1985 | 0.6434 | 0.8763 | 1.46E-43 | 0.4284 |
|  | Average accessible surface area | Janin et al., 1978 | 22.1445 | 30.9438 | 3.21E-42 | 0.4214 |
| **Heterologous Interfaces** | Normalized positional residue frequency at helix termini C' | Aurora-Rose, 1998 | 0.4753 | 0.7315 | 5.83E-37 | 0.4343 |
|  | Side chain angle theta(AAR) | Levitt, 1976 | 19.9685 | 36.1640 | 6.45E-34 | 0.4149 |
|  | Normalized frequency of coil | Tanaka-Scheraga, 1977 | 0.4166 | 0.6041 | 2.73E-33 | 0.4108 |
| **Heteromeric Interfaces** | Principle property value z3 | Wold et al., 1987 | 1.9069 | 2.5636 | 1.36E-72 | 0.4014 |
|  | Average non-bonded energy per atom | Oobatake-Ooi, 1997 | 0.1720 | 0.2293 | 5.64E-71 | 0.3968 |
|  | Apparent partition energies calculated from Janin index | Guy, 1985 | 0.6333 | 0.8459 | 7.29E-71 | 0.3965 |
| **DNA interfaces** | Dependence of partition coefficient on ionic strength | Zaslavsky et al., 1982 | 0.0901 | 0.1288 | 3.37E-12 | 0.4153 |
|  | Normalized frequency of N-terminal helix | Chou-Fasman, 1978 | 0.4458 | 0.6259 | 9.71E-11 | 0.3857 |
|  | Helix termination parameter at position j+1 | Finkelstein et al., 1991 | 0.3895 | 0.5264 | 3.78E-09 | 0.3510 |
| **RNA interfaces** | Free energy change of $\alpha(R_i)$ to $\alpha(R_h)$ | Wert-Scheraga, 1978 | 0.3104 | 0.4626 | 4.54E-05 | 0.5604 |
|  | Partial specific volume | Cohn-Edsall, 1943 | 0.0855 | 0.1148 | 3.18E-04 | 0.4945 |
|  | Hydrophobicity coefficient in RP-HPLC, C18 with 0.1%TFA/MeCN/H<sub>2</sub>O | Wilce et al. 1995 | 1.5660 | 2.0809 | 3.64E-04 | 0.4897 |
| **Ligand interfaces** | Average non-bonded energy per atom | Oobatake-Ooi, 1977 | 0.1766 | 0.2235 | 1.29E-60 | 0.3193 |
|  | Percentage of exposed residues | Janin et al., 1978 | 19.8774 | 25.2315 | 3.13E-58 | 0.3127 |
|  | Apparent partition energies calculated from Janin index | Guy, 1985 | 0.6498 | 0.8204 | 4.74E-58 | 0.3122 |

There are a several properties with high effect sizes that distinguish pathogenic from benign mutants at multiple locations. These include mutants that affect hydrophobicity, hydrophilicity and related properties such as percentage of buried residues and accessible surface area. Since an altered hydrophobic profile can influence protein folding, this is unsurprising. *Average non-bonded energy per atom*, the property with the largest effect size over all locations, is the calculated contribution of each residue to the non-bonded energy of proteins. This property is also related to hydrophobicity, as hydrophobic interactions contribute greatly towards the total non-bonded energy (46). *Principle property value z3* (47) is also among the top properties that distinguish pathogenic from putatively benign mutations at interior and surface locations as well as heteromeric interfaces. This is essentially a principal component derived from multiple other properties.

Most of the properties with the highest effect sizes are either related to hydrophobicity or to free energy changes, with the exception of two locations: homomeric heterologous interfaces and DNA interfaces. In these locations, properties related to secondary structure, helices specifically, also have moderate effect sizes. This hints at potentially different mechanisms of pathogenesis at each of these sites via the perturbation of secondary structure.

The change in effect size ranking for each property was calculated for every pair of locations. We visualised the difference in the property changes in Figure 6. These graphs plot change in rank against change in effect size for all AAindex properties at a pair of locations. A wider distribution indicates greater differences between the properties changes occurring in pathogenic vs benign mutations at the two locations. Properties that are highly skewed towards one corner of the plot show a large pathogenic/benign property difference at one location, but a small one at the other, indicating that these properties may be considerably more important for residues at one location than the other. Also marked on the plots are those properties with the greatest effect size at each location. The presence of these properties at the centre of the plot indicates that these properties are equally important at the two locations, and if they are highly skewed, then the opposite.

**Figure 6:**
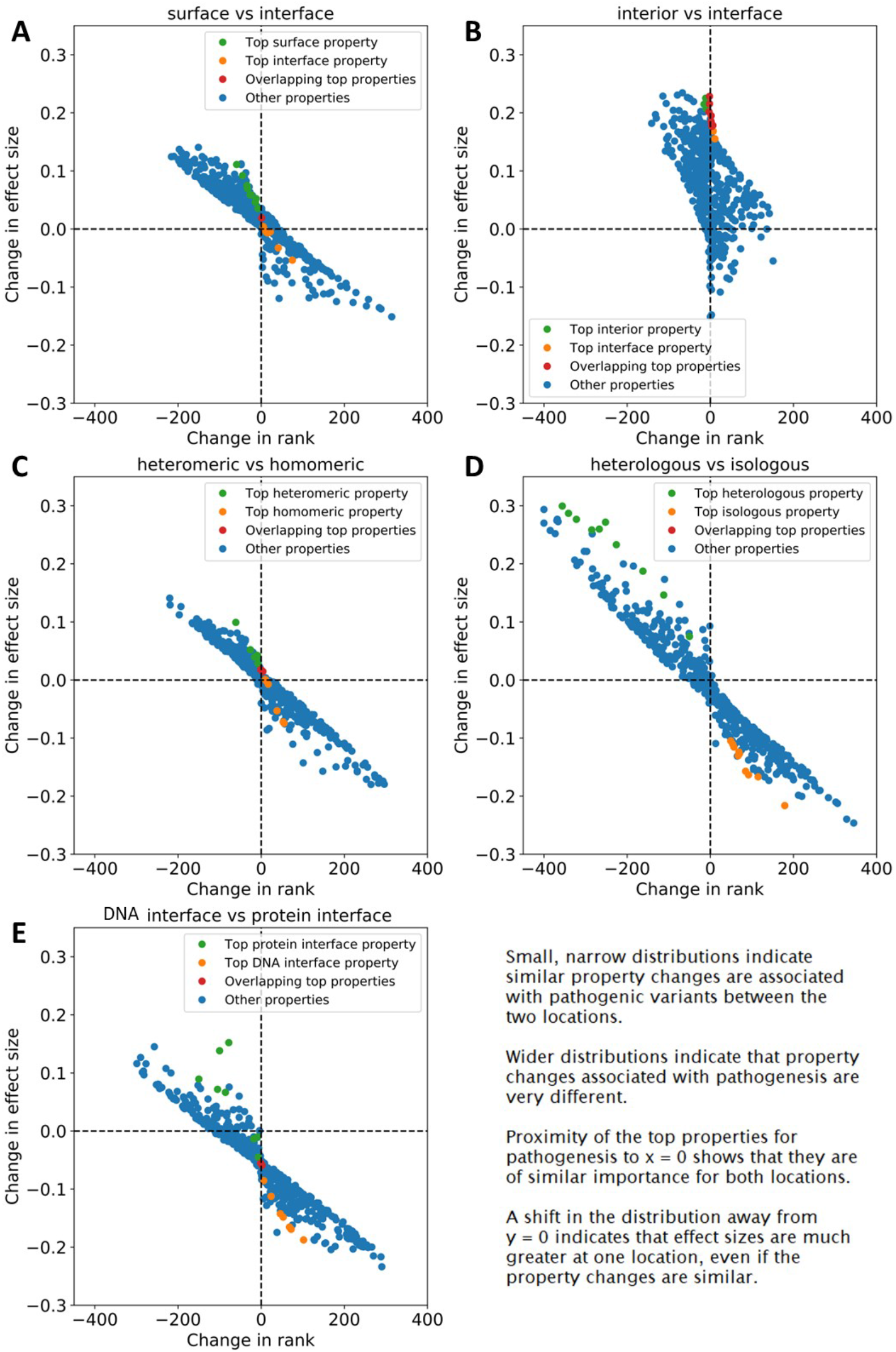
Differences in property changes (delta properties) between locations plotted against rank change of those properties. Effect size difference in the delta properties of two locations is on the Y-axis, and rank change of the delta properties is on the X-axis. The top properties (by effect size) for each location are coloured green and orange, while overlapping top properties are coloured red. A) Surface locations vs all interfaces. B) Interior locations vs all interfaces. C) Heteromeric vs homomeric interfaces. D) Heterologous vs isologous interfaces. E) DNA interfaces vs all protein interfaces.

To address the question of whether interfaces resemble interior or surface locations the most, we plotted surface vs interface property changes (Fig 6A) and interior vs interface property changes (Fig 6B). The wider distribution on the x-axis in Fig 6A demonstrates that the property changes important for pathogenesis are more different between surface vs interface positions, and more similar between interior vs interface positions. This is further corroborated by the fact that the top properties associated with pathogenesis in each location do not overlap in Fig 6A, whereas there is considerable overlap between the top properties in Fig 6B, indicating that the most important properties for pathogenesis are the same at both locations. In addition, the entire plot in Fig 6B is shifted towards positive y-axis values, due to increased effect sizes of these properties at interior locations, despite similar relative importance at each location.

In heteromeric vs homomeric (isologous and heterologous combined) interfaces (Fig 6C), the distribution is slightly more compact than in Fig 6A. The limited overlap between top properties indicates that more similar property changes are responsible for pathogenesis at these locations than between surface and interface amino acids. However, when we compare heterologous interfaces to isologous (Fig 6D) we find a much wider distribution and complete separation of the top properties. The top properties in this distribution are among those furthest from the origin and are thus those that differ in effect on pathogenesis the most between the two locations. These were primarily properties related to secondary structure such as *relative preference value at C-cap* (48) and *normalized frequency of left-handed alpha-helix* (49). This result could be biased by the presence of transmembrane helices in cyclic membrane proteins such as ion channels; however, exclusion of predicted membrane proteins yielded similar results (Fig S4).

DNA interfaces also showed considerable differences when compared to protein-protein interfaces (Fig 6E). Although there was a single overlap between the top properties, they also represent some of those furthest from the origin, particularly *normalized frequency of N-terminal helix* (50) and *dependence of partition coefficient on ionic strength* (51). Secondary structure once again appears to be a major factor. It is likely that the major differences between DNA interfaces and other interface types are those mutations that alter DNA specificity or abolish DNA binding.

### Performance of variant effect predictors on interface mutations

One of the areas of biology which has most benefited from the recent explosion in genome sequencing and increase in understanding of the mechanisms of pathogenesis, is that of variant effect prediction (52–54). The effect of most coding variants on phenotype remains completely unknown; therefore, these predictors are valuable in both clinical and research environments. The basis of almost all predictors is evolutionary conservation; however, the inclusion of additional features also often positively contributes towards accurate predictions. Given the varying properties of disease mutations at different locations we observed in the previous section, we believe that this could have implications for variant effect predictions.

In order to assess the current state of computational variant effect prediction by protein region, we obtained results from 29 computational predictors of variant effect from the dbNSFP database (55). We then retrieved predictions for as many mutations in our datasets as possible and calculated the receiver operating characteristic, area under the curve (ROC AUC) statistic for each interface type and location for each predictor. The calculations were repeated 1000 times by resampling the pathogenic and putatively benign datasets with replacement, and the average AUC statistic was taken. Regardless of the predictor methodology, there are several common patterns. The results for four representative predictors (SIFT: an empirical method (56), DEOGEN2: a supervised method (57), PROVEAN: an unsupervised method (58) and REVEL: a metapredictor (59)) are shown in Figure 7, while full results are provided in Figure S5.

**Figure 7:**
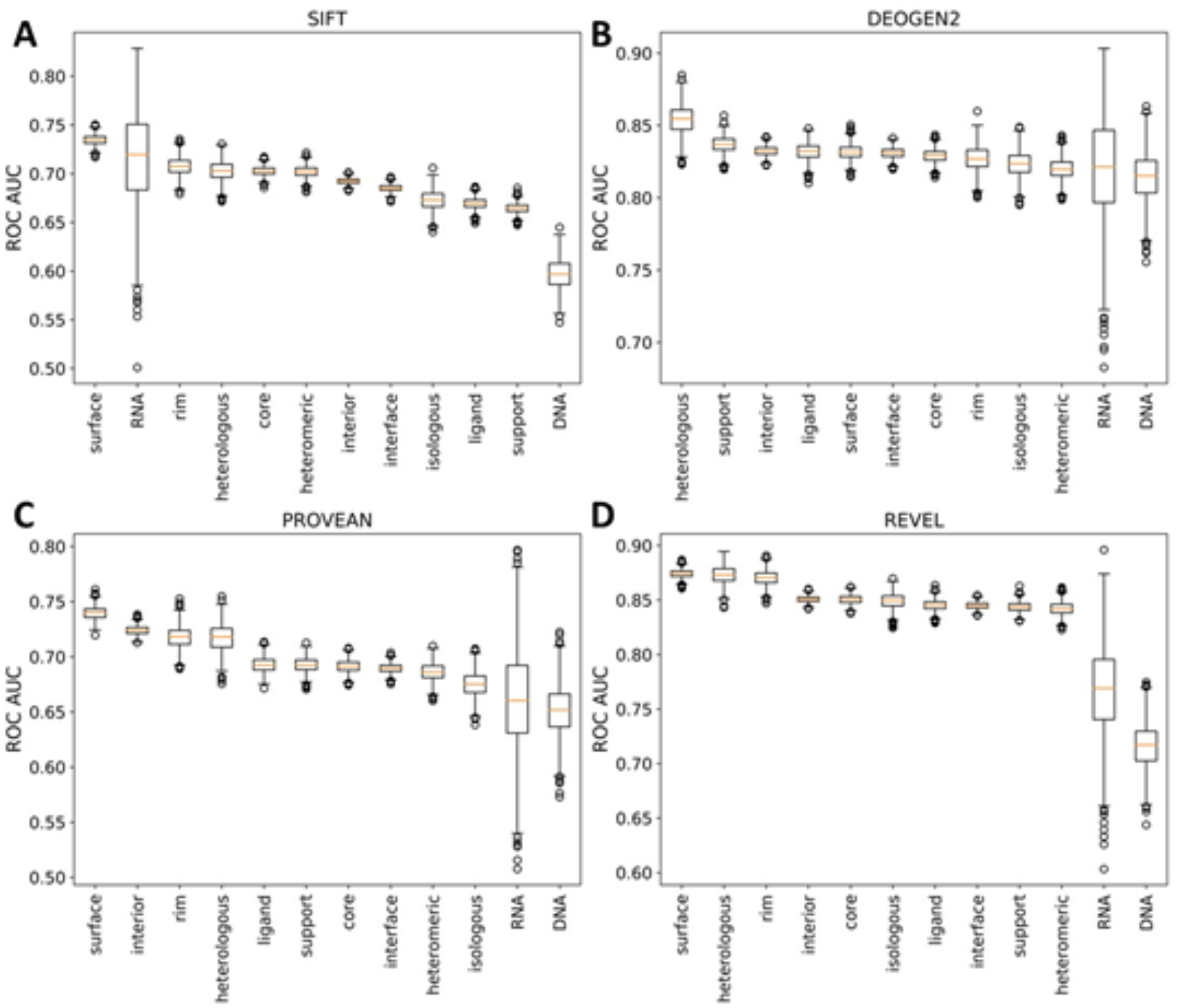
Distribution of bootstrapped ROC AUC for predictions made by four representative VEPs at different protein locations. A) SIFT: an empirical predictor. B) DEOGEN2: a supervised machine learning method. C) PROVEAN: an unsupervised machine learning method. D) REVEL: a metapredictor (*i*.*e*. ensemble predictor). Charts for all predictors are available in figure S5.

Some interesting patterns emerge when comparing the performance of variant effect predictors across different protein locations. Out of surface, interior and interface mutations, surface mutants are most frequently the best predicted, while interfaces are the worst, a trend that matches their relative levels of pathogenic mutations. Between the different interface types, the outcomes of mutations at heterologous interfaces tend to be predicted particularly well, being better predicted than those at isologous interfaces for 28/29 predictors, and better than any interface locations for 18 predictors. In contrast, pathogenic mutations at DNA interfaces tended to be predicted quite poorly, being the worst predicted interface type for 21 predictors.

Of all the predictors used, only DEOGEN2 (Fig 7B) specifically includes protein-protein interface information as a feature. However, this did not appear to have a great impact on its ROC statistic, as interfaces overall were still worse predicted than interior or surface locations. DEOGEN2 also still underperforms REVEL (Fig 7D) at all locations except DNA and RNA interfaces.

The consistency at which mutations at certain interface types are better predicted than others by variant effect predictors further demonstrates that the interface type has an impact on prediction efficiency across multiple methods. The consistency at which heterologous interfaces were better predicted than isologous interfaces was surprising. One possible explanation for this could be that complexes with heterologous interfaces tend to be more conserved and thus able to be better predicted than homodimers. The mechanisms of DNA and RNA binding, on the other hand are unlikely to be effectively captured by non-specialist variant effect predictors, as related transcription factors can bind to very different sequences.

## Conclusions

The goals of this study have been to assess the nature of differences in pathogenic and benign mutations in the context of protein interfaces, with a specific focus on findings that may inform or benefit future variant effect prediction methodology. The division of protein interfaces based on ligand type and on overall complex symmetry is an approach that has received little previous attention. Heterologous protein interfaces in particular demonstrated an increased enrichment of disease mutations relative to other protein-protein interface types. This was further demonstrated when cyclic, dihedral and helical complexes (all of which can contain heterologous interfaces) were also found to be enriched in pathogenic mutations. We ruled out ion-channel enrichment as a possibility for this observation. A more likely scenario is that, since heterologous interfaces are a basis for the generation of larger complexes (through formation of fibrils, or larger closed cycles), and that slightly disruptive interface mutations have a larger effect on complex stability than on the stability of smaller complexes such as dimers. It is also likely that this effect increases with the number of mutant subunits forming the protein complex.

All locations within the protein demonstrated broadly similar patterns of tolerance towards specific amino acid substitutions. For example, regardless of location, mutations away from cysteine were damaging, as were mutations to proline. However, comparison of the relative enrichment ratios between locations pointed towards some interesting relationships. Mutations from cysteine on the protein surface were found to be relatively more damaging than at all other locations. Previous work has suggested that there is selection pressure to remove cysteines from the protein surface due to its high reactivity (60); this implies that remaining surface cysteines have increased functional importance, thus explaining our observations. We also found evidence of differences between amino acids involved in pathogenic mutations within isologous and heterologous interfaces, and within DNA and protein interfaces.

Allostery has been of great interest to the scientific community of late, particularly in the field of drug discovery (61,62). Our observation that sites distal to protein interfaces are enriched in pathogenic mutations in a distance-dependent manner is most likely a form of allostery. Indeed, allostery has been implicated as a mechanism involved in several disease mutations (40,41).

These findings also have implications for future variant annotations (both manual and through use of computational tools). While 3D protein structural information has been integrated into a number of variant effect predictors (63,64), interface proximity has not, so far, been used as a predictive feature, to the best of our knowledge. Our results indicate that inclusion of interface proximity as a feature may be beneficial in predicting variant pathogenicity.

Heterologous homomeric protein-protein interfaces have demonstrated some unexpected characteristics in comparison to other protein-protein interface types, even in comparison to isologous homomeric interfaces. This is most greatly reflected in the properties that exhibit the most change between pathogenic and benign mutations. While changes in hydrophobicity and related properties are among the most important for pathogenesis at almost every location, for heterologous interfaces (along with DNA interfaces), properties relating to secondary structure exhibit greater effects. While it is tempting to explain this as being due to transmembrane helices in membrane ion channels (of cyclic symmetry), removal of these proteins from the analysis does not alter the observation. It may be that secondary structure disruption is a more important factor at heterologous interfaces than altered hydrophobicity, which is an important observation for variant effect prediction at these sites. The case for protein-DNA interfaces is easier to explain, since DNA binding motifs often involve alpha helical motifs (65).

Computational variant effect predictors have been widely used for decades now. These programs are most commonly based on evolutionary conservation within a multiple sequence alignment. As a proxy metric for mutation tolerance, it performs well and is a major part of even the most modern predictors. Additional biochemical and structural features are also often included where they can aid performance. One aspect that is rarely considered when training a predictor, is the presence or the absence of a protein interface, or the type of protein interface. The consistent bias of all predictors in favour of heterologous interfaces and against DNA/RNA interfaces underscores the need for improvement in this area. Our results indicate several possible features which may be of benefit to future variant effect predictors including interface presence and type, distance to interface and location-specific property changes.

Our work has highlighted a number of interesting aspects of pathogenic mutations at protein interfaces. We hope that these findings will help improve variant effect predictor methodology and inspire further research in the area.

## Materials and Methods

### ClinVar and gnomAD databases

To establish sets of pathogenic and benign mutations, we used the ClinVar (28), downloaded on 2020.08.17, and gnomAD v2.1.1 (29) databases. After filtering for missense variants, we selected only those variants in ClinVar labelled as “pathogenic” and “likely pathogenic”. Variants in gnomAD are aggregated from large-scale sequencing projects and exclude those individuals with severe paediatric disease. We removed all variants shared between both datasets from the gnomAD set.

### Protein structures

Protein structures were downloaded from the Protein Data Bank (PDB) (66) on 2020.05.27. The first biological assembly was used to represent the quaternary structure of each protein, and symmetry assignments were taken directly from the PDB. We searched for those polypeptide chains with >90% sequence identity to a human protein over a region of at least 50 amino acid residues. For human protein residues that map to multiple chains, we selected a single chain sorting by highest resolution followed by largest structure. The full set of human ClinVar and gnomAD mutations mapped to protein structures is provided as a supplementary dataset.

### SASA calculations

SASA was calculated for each structure using AREAIMOL (67). Residues were determined to be members of a protein interface if they experienced a change in SASA between the monomeric structure and the structure of the full PDB complex. SASA was then scaled (30) to determine the fraction of exposed residue. Residues with greater than or equal to 25% exposed surface area were classified as surface residues, while those with less were classified as interior. Those residues that changed from surface to interior upon complex formation were classified as interface core residues. Those that experienced a change in SASA but remained interior were classified as support residues and those that experienced a change but remained surface were classified as rim residues (Fig 1A).

### Odds ratios

Odds ratios, or relative enrichment values were calculated using the scipy.stats.fisher_exact() function from the python scipy package. For a pathogenic and putatively benign group within a subset of the data, this is equivalent to:

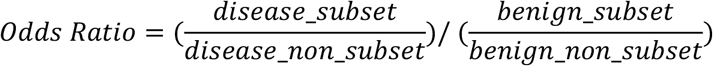

Confidence intervals (95%) for odds ratios were determined by:

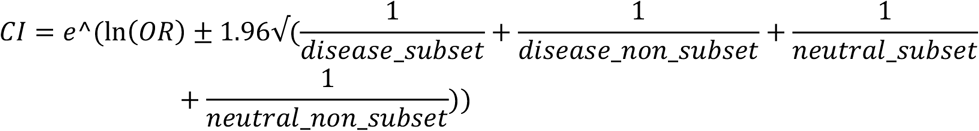

where CI is the upper/lower confidence interval and OR is the odds ratio.

### 3D complex residue distance calculations

Distances between interior/surface mutations and interfaces were calculated using atomic coordinates in PDB files. The coordinates of alpha-carbon atoms were used to represent the location of all residues. Rim residues were removed from all interfaces. For each PDB structure, the centroid of each interface was calculated as the mean of the x, y and z coordinates of every amino acid in the interface. The distance between each interior/surface mutation and interface was calculated using:

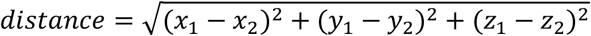

for each interface within the PDB chain. Where multiple interfaces were present in the structure, then the distance to the closest interface was used to represent the proximity of the mutation. To calculate linear sequence proximity to an interface, the absolute sequence distance to the closest interface residue was used.

### Variant effect predictors

Variant effect predictions were obtained from the dbNSFP database (55), with the exception of SIFT which was run locally using the Uniref90 database (68) to generate alignments.

### ROC curves

Receiver operating characteristic (ROC) area under the curve (AUC) statistics were generated using the sklearn.metrics.roc_auc_score() function of the sklearn python package. Several VEPs used inverse metrics to score variants (lower, rather than higher score representing greater probability of damaging variants). To resolve this issue, we deducted the score from 1.0 for all VEPs with an AUC less than 0.5.

### AAindex

Amino acid properties and substitution matrices were obtained from the AAindex database (45). Properties without values for all 20 standard amino acids were removed from the analysis.

### Property deltas

Absolute differences in properties were calculated for individual amino acid substitutions. A two-tailed Student’s t-test was then performed to determine significance of the difference between the mean of pathogenic and putatively benign mutation properties at different locations. Hedges’ g was used to calculate effect size.

## Data availability

All datasets related to this study are available at https://doi.org/10.7488/ds/3116

## Acknowledgements

This work was supported by an MRC Career Development Award (MR/M02122X/1) to JM and an MRC Precision Medicine Doctoral Training Programme studentship to BL. JM is a Lister Institute Research Prize Fellow.

## Figures

**Figure S1:**
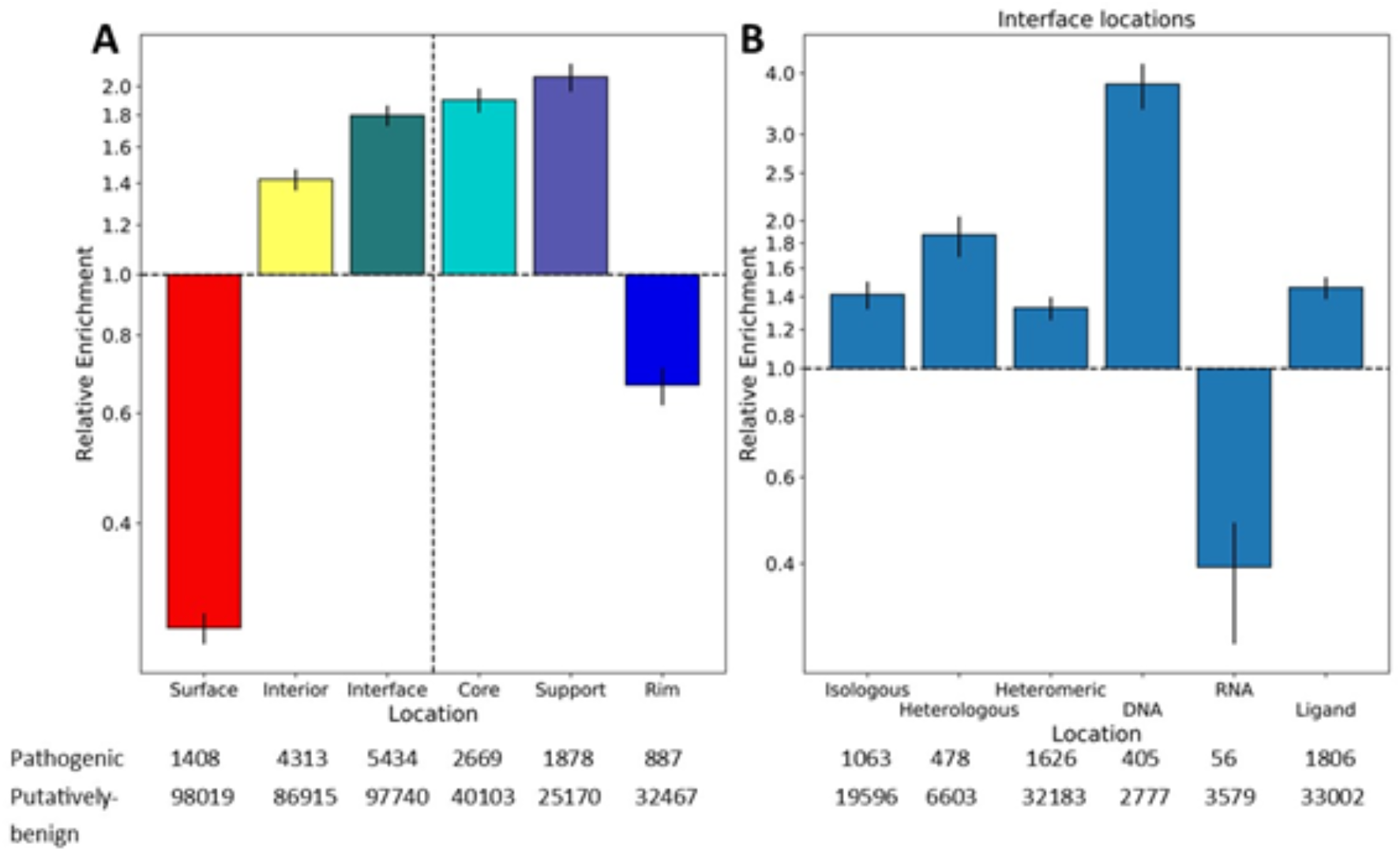
Enrichment of pathogenic variants at different protein locations and interface types excluding predicted membrane proteins. Error bars represent 95% confidence intervals. A) Enrichment of pathogenic variants in surface, interior and interface locations relative to all data. B) Enrichment of pathogenic variants in different interface types relative to all data. Total number of pathogenic and putatively benign variants is shown below the plots.

**Figure S2:**
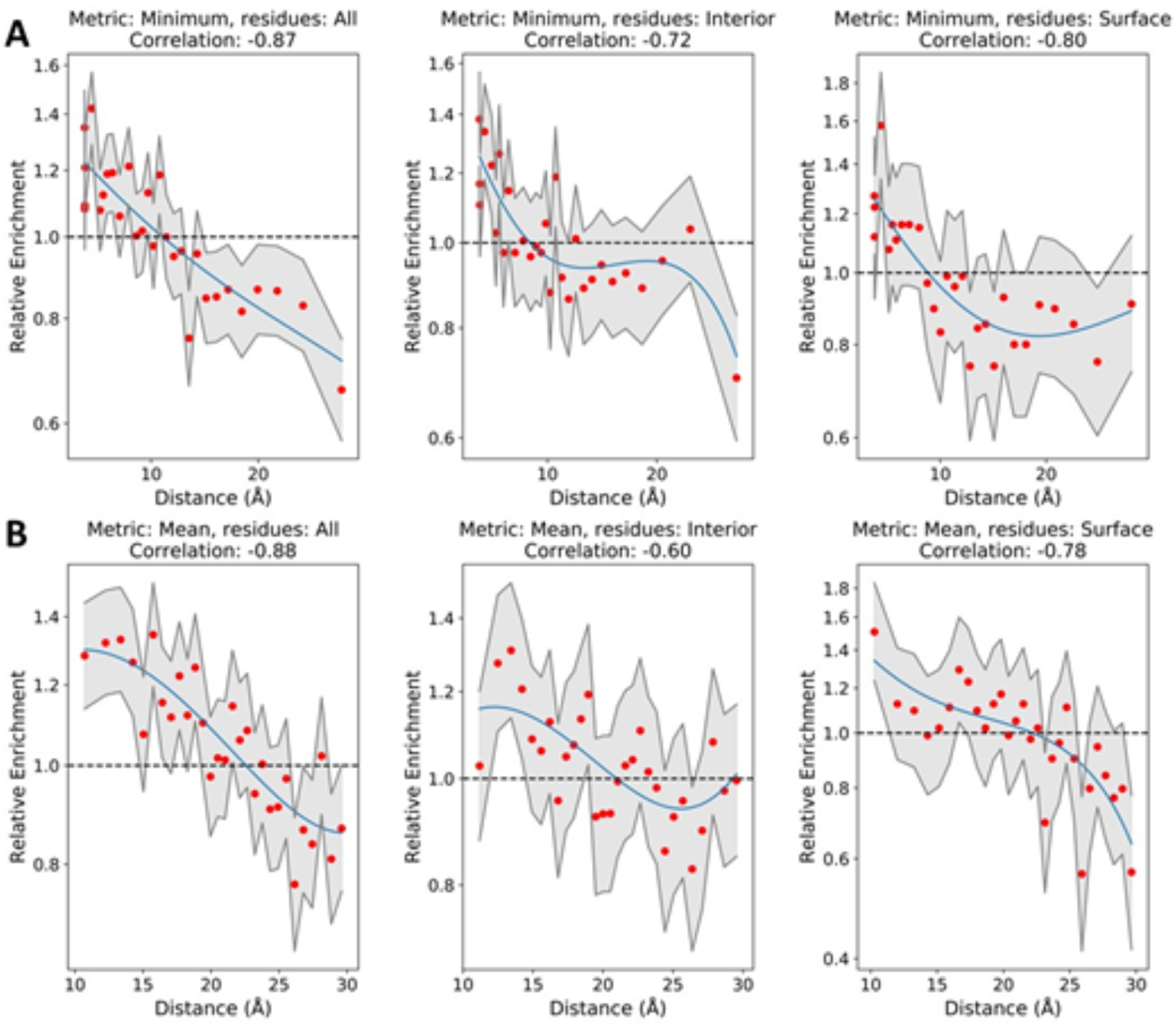
Log odds ratio of disease mutations at increasing distance from the nearest interface. A) Using the minimum distance to the nearest interface residue. B) Using the mean distance to all interface residues. 95% confidence intervals are shown in grey, a univariate spline (blue) has been fit to the data. Charts are shown for surface and interior residues together and individually.

**Figure S3:**
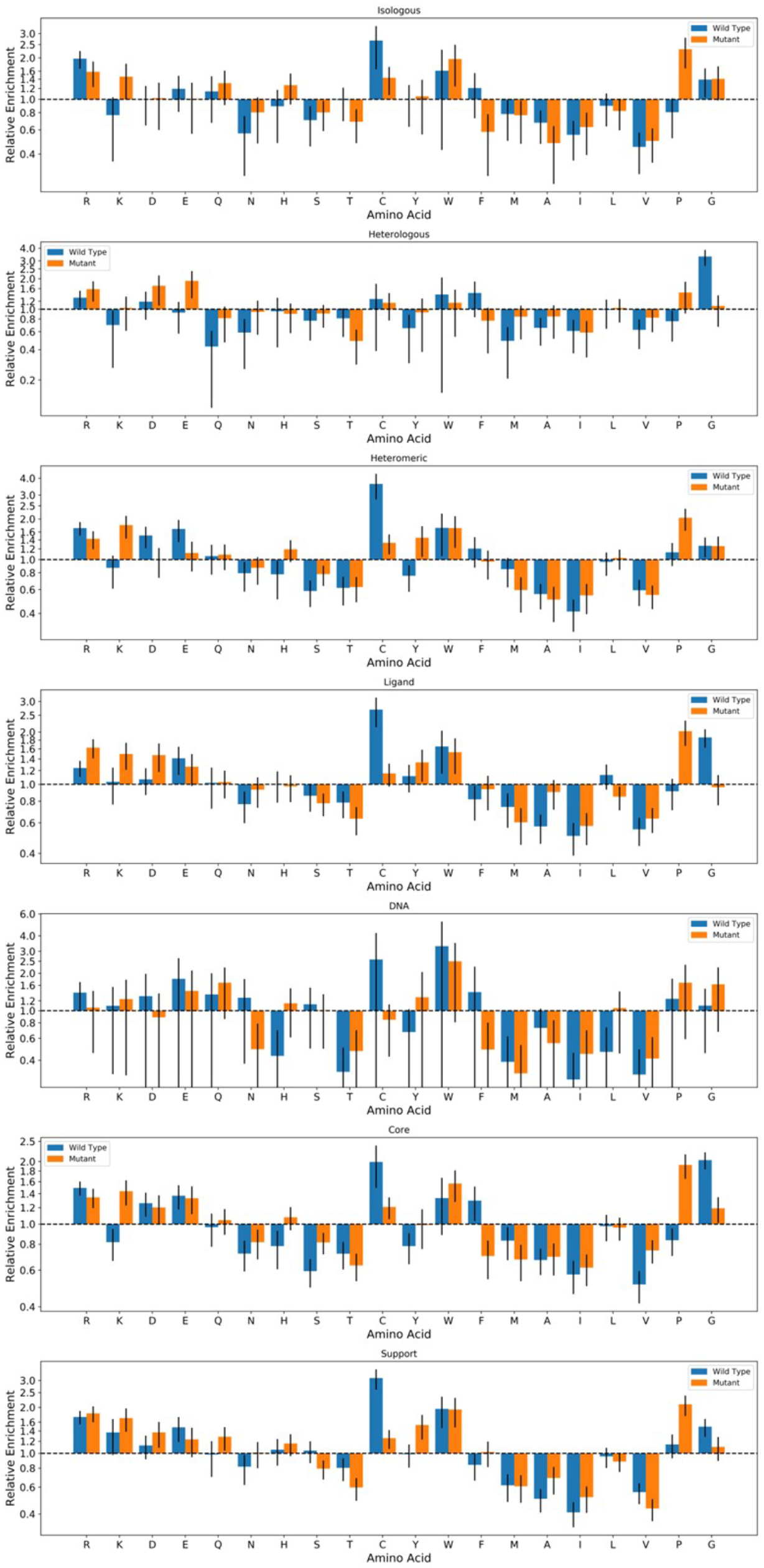
Enrichment of pathogenic variants at specific residues at different interface types. The log odds ratio of pathogenic mutations associated with specific amino acid mutations at each interface type as well as core and support regions. Blue bars represent mutations *from* a specific amino acid, while orange bars represent mutations *to* a specific amino acid. All error bars represent 95% confidence intervals.

**Figure S4:**
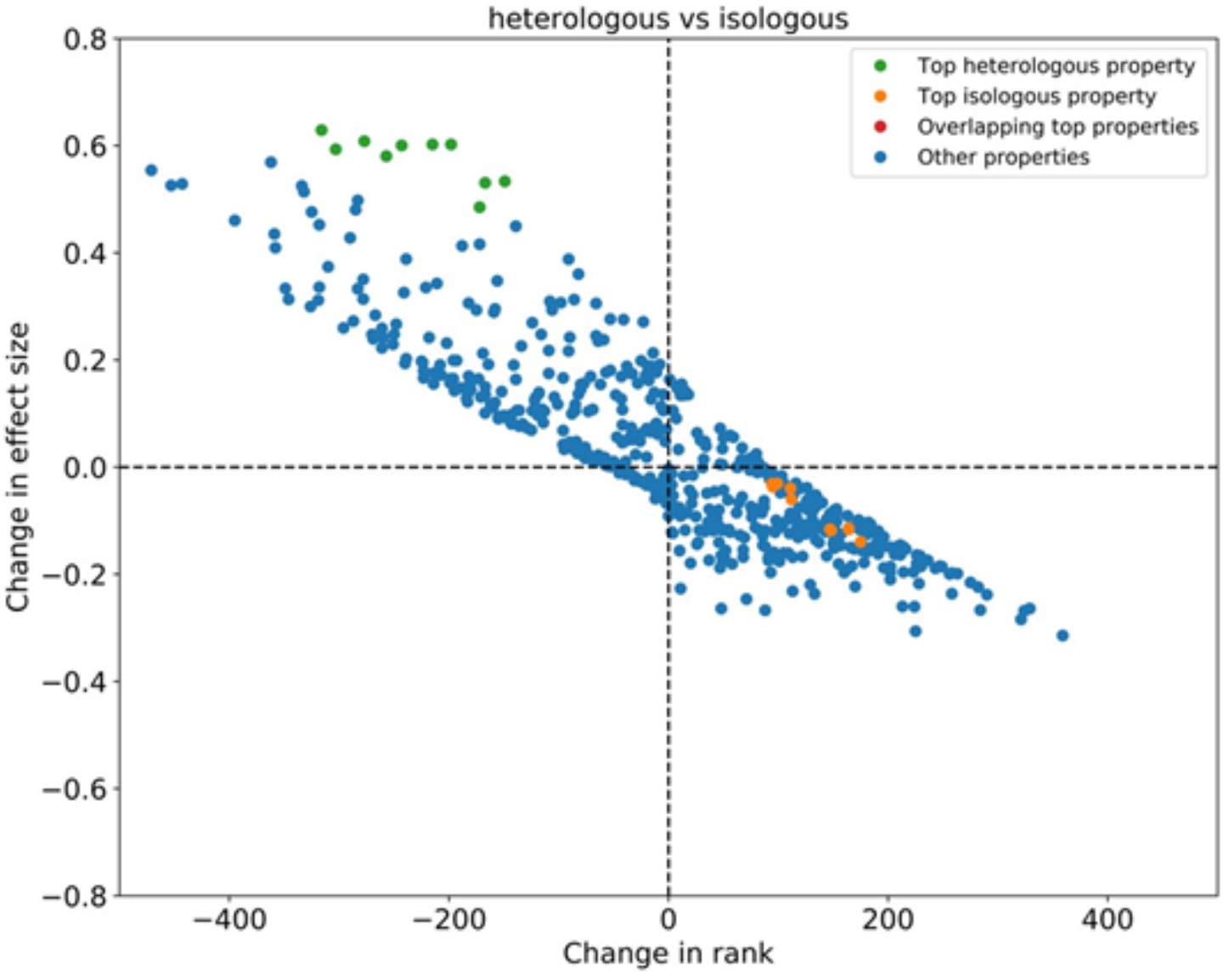
Difference in property changes between isologous and heterologous interfaces plotted against rank change of those properties with all predicted membrane proteins excluded. Effect size difference in the delta properties of two locations is on the Y-axis, and rank change of the delta properties is on the X-axis. The top properties (by effect size) for each location are coloured green and orange.

**Figure S5:**
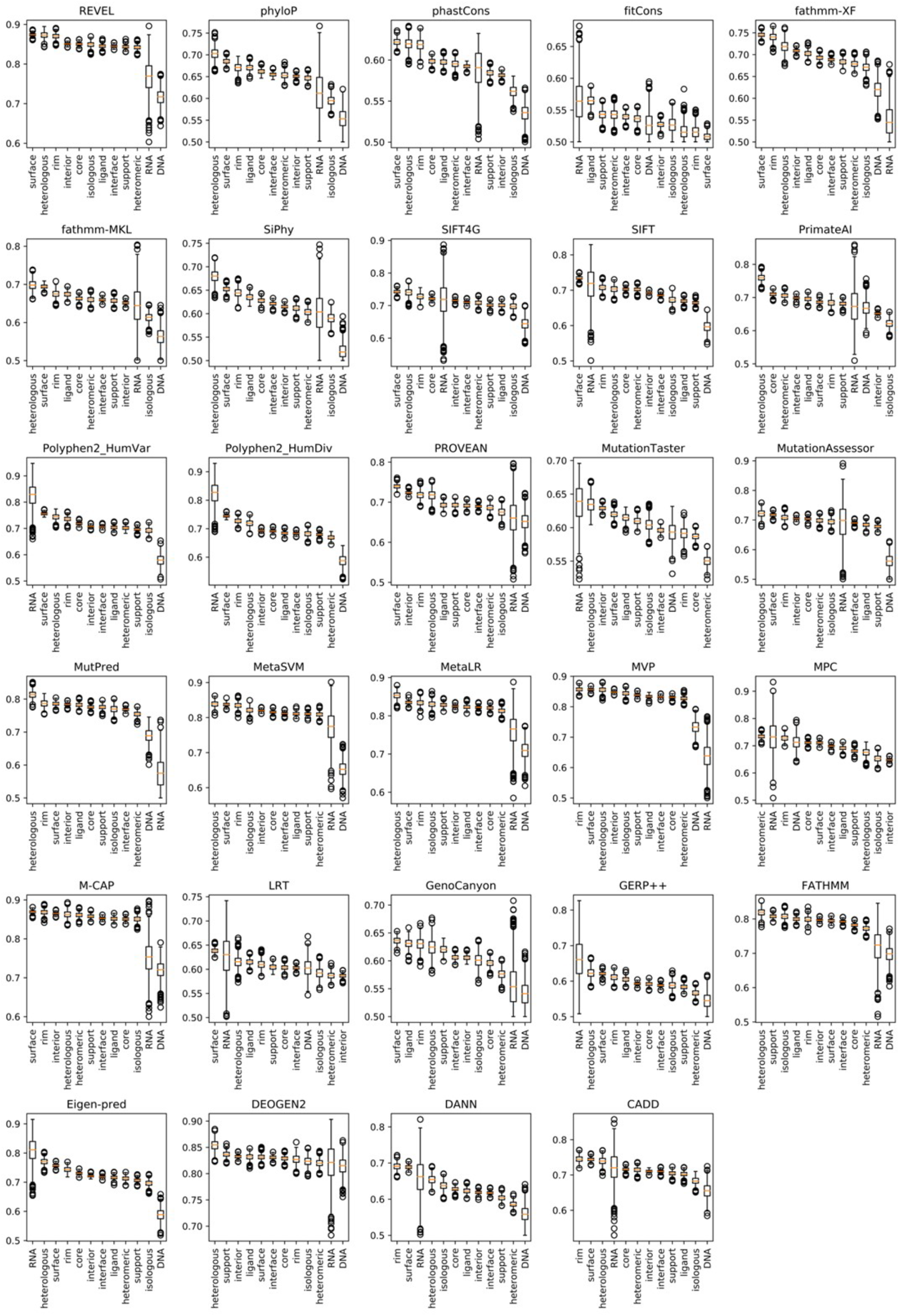
Distribution of bootstrapped ROC AUC for predictions made by VEPs at different protein locations. Pathogenic and putatively benign datasets were independently re-sampled 1000 times with replacement. Only mutations with all 29 predictions were included in this analysis.

